# Using ultraconserved elements to reconstruct the termite tree of life

**DOI:** 10.1101/2021.12.09.472027

**Authors:** Simon Hellemans, Menglin Wang, Nonno Hasegawa, Jan Šobotník, Rudolf H. Scheffrahn, Thomas Bourguignon

**Affiliations:** Okinawa Institute of Science & Technology Graduate University, 1919-1 Tancha, Onna-son, Okinawa 904-0495, Japan; Faculty of Tropical AgriScience, Czech University of Life Sciences, Kamýcka 129, 16521 Prague, Czech Republic; Fort Lauderdale Research and Education Center, Institute for Food and Agricultural Sciences, 3205 College Avenue, Davie, Florida 33314 USA

**Keywords:** Data Mining, Isoptera, Phylogenomics, Mitochondrial genome, Nuclear genome, Taxonomy

## Abstract

The phylogenetic history of termites has been investigated using mitochondrial genomes and transcriptomes. However, both sets of markers have specific limitations. Mitochondrial genomes represent a single genetic marker likely to yield phylogenetic trees presenting incongruences with species trees, and transcriptomes can only be obtained from well-preserved samples. In contrast, ultraconserved elements (UCEs) include a great many independent markers that can be retrieved from poorly preserved samples. Here, we designed termite-specific baits targeting 50,616 UCE loci. We tested our UCE bait set on 42 samples of termites and three samples of *Cryptocercus*, for which we generated low-coverage highly-fragmented genome assemblies and successfully extracted *in silico* between 3,426 to 42,860 non-duplicated UCEs per sample. Our maximum likelihood phylogenetic tree, reconstructed using the 5,934 UCE loci retrieved from upward of 75% of samples, was congruent with transcriptome-based phylogenies, demonstrating that our UCE bait set is reliable and phylogenetically informative. Combined with non-destructive DNA extraction protocols, our UCE bait set provides the tool needed to carry out a global taxonomic revision of termites based on poorly preserved specimens such as old museum samples. The Termite UCE database is maintained at: https://github.com/oist/TER-UCE-DB/.

## 1. Introduction

Termites are the most ancient lineage of social insects, with a fossil record dating back to the Early Cretaceous, ∼135 million years ago (Mya) (Thorne *et al*., 2000; Grimaldi & Engel, 2005; Engel *et al*., 2009). All modern termites share a common ancestor estimated at 140-150 Mya by time-calibrated phylogenetic trees (Bourguignon *et al*., 2015; Bucek *et al*., 2019). However, the bulk of the modern termite species diversity belongs to the Termitidae, a lineage that originated during the early Eocene, ∼50 Mya, as indicated by time-calibrated phylogenetic trees (Bourguignon *et al*., 2015, 2017; Bucek *et al*., 2019) and by the fossil record (Engel *et al*., 2011). While the backbone of the phylogenetic tree of termites is now largely resolved, most termite species are still awaiting to be placed on the tree of life.

Our understanding of the phylogenetic history of termites was mostly based on mitochondrial markers and fossils to calibrate estimated times of divergence until Bucek *et al*. (2019) published a phylogenetic tree of termites based on transcriptome data. The first phylogenetic trees of termites were based on a couple of PCR-amplified mitochondrial markers, sometimes combined with nuclear 18S or 28S sequences and/or morphological characters, which hardly contributed any phylogenetic signal (e.g., Lo *et al*., 2004; Inward *et al*., 2007; Legendre *et al*., 2008). These phylogenies provided a good overview of the relationships among the main termite lineages but lacked the robustness of phylogenetic trees inferred from full mitochondrial genomes (e.g., Cameron *et al*., 2012; Bourguignon *et al*., 2015, 2017). Full mitochondrial genomes, which became easy to sequence with the rise of second-generation sequencing technologies, resolve both shallow and deep divergences in the evolutionary history of termites and other insect lineages (Cameron, 2014), making them a marker of choice for phylogenetic reconstructions. However, mitochondrial genomes form a single marker, as all mitochondrial genes are linked and maternally inherited as a single package. Consequently, mitochondrial phylogenies are sometimes discordant with species phylogenies, especially for closely related species and short internal branches that diverged in periods of time too brief for alleles to coalesce (Whitfield & Lockhart, 2007; Degnan & Rosenberg, 2009). One example of such discordance is provided by Sphaerotermitinae, the unambiguous sister group of Macrotermitinae according to transcriptomic data; but supported as sister to non-macrotermitine non-foraminitermitine Termitidae by mitochondrial genome phylogenies (Figure 1; Bucek *et al*., 2019). Phylogenies based on multiple independent nuclear markers are needed to resolve the evolutionary history of organisms accurately.

**Figure 1:**
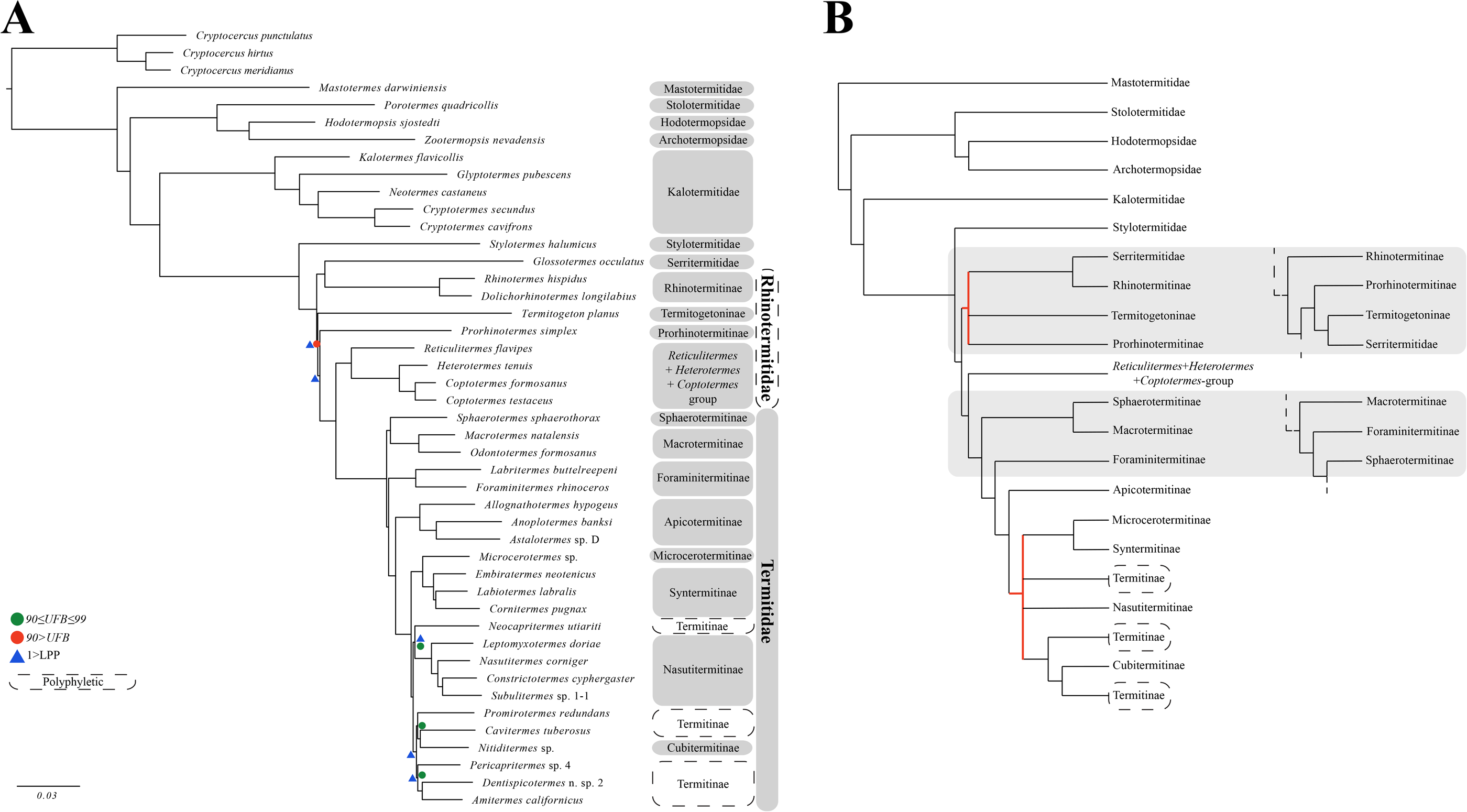
(**A**) Maximum likelihood phylogenetic tree of termites reconstructed with IQ-TREE using 5,934 UCE loci and complete mitochondrial genomes. Only UCE loci present in more than 75% of species were used. Support values are indicated for non-fully resolved nodes: ultrafast bootstrap (UFB; summarized from the phylogenetic trees reconstructed using UCE only and UCE + mitochondrial DNA displayed in Figures S3 and S4, respectively) and ASTRAL-III local posterior probabilities (LPP; phylogenetic tree displayed in Figure S5) values. (**B**) Family-level summary topology of termites supported by both UCEs (this study) and transcriptomic data (Bucek *et al*., 2019), with the indication of alternative topologies inferred from mitochondrial genome data alone (Bourguignon *et al*., 2015, 2017). Unsupported splits were summarized as polytomies (branches in red).

Transcriptomes, the snapshot of genes expressed by an organism during tissue sampling, include many independent nuclear markers that can be used to build robust phylogenetic trees. Transcriptome-based phylogenies, reconstructed using up to ∼4,000 single-copy orthologous nuclear genes spanning over 7.7 million nucleotide positions, have provided a robust picture of the ancient evolutionary history of termites (Bucek *et al*., 2019). The sequencing of transcriptomes is now affordable, but, unfortunately, RNA is unstable and can only be extracted from samples that have been adequately preserved and stored, preventing the use of most samples collected before the genomic era began and making the approach impractical for large-scale studies. One alternative is to mine the conserved genetic markers present in whole-genome shotgun sequencing datasets, such as some datasets generated to sequence mitochondrial genomes.

Ultraconserved Elements (UCEs) are highly conserved nuclear regions whose functions remain largely unknown (Bejerano *et al*., 2004; Faircloth *et al*., 2012). UCEs are found across all regions of animal genomes, including the exonic, intronic, and intergenic regions. Phylogenetic trees inferred from UCEs have contributed to our understanding of the evolutionary history of various animal lineages spanning across the animal tree of life (e.g., Faircloth *et al*., 2012; Ryu *et al*., 2012; Smith *et al*., 2014; White & Braun, 2019; Zhang *et al*., 2019). Unlike transcriptomes, UCEs can readily be obtained from museum samples through baiting conserved elements and their phylogenetically-informative flanking regions from fragmented genome assemblies (Blaimer *et al*., 2016; Faircloth, 2017; Derkarabetian *et al*., 2019). No UCE bait set has been designed for termites so far. We filled this gap as follows: (*i*) we designed *in silico* baits to capture UCEs; (*ii*) we compared phylogenetic trees reconstructed using all possible combinations of mitochondrial genomes, nuclear ribosomal RNA genes, and UCEs; and (*iii*) we showed that UCEs obtained from low-coverage shotgun genome assemblies are an alternative to transcriptome-based phylogenies for the reconstruction of multi-gene phylogenies. Finally, we set up a Termite UCE Database, thereby ensuring a long-term re-usability of published data.

## 2. Material and Methods

### Biological samples and sequencing

We used sequence data from 42 samples of termites and three samples of *Cryptocercus*, the wood-feeding subsocial cockroach genus forming the sister group of termites. The sequencing data of 14 species were retrieved from previous studies (for details, see Table S1). The sequencing data from the remaining 31 species were obtained from samples preserved in 80% ethanol stored at room temperature or from samples preserved in RNA-later® and stored at temperatures ranging between -20°C and -80°C until DNA extraction. DNA was extracted using the DNeasy Blood & Tissue extraction kit (Qiagen). Libraries were prepared using the NEBNext® Ultra™ II FS DNA Library Preparation Kit (New England Biolabs) and the Unique Dual Indexing Kit (New England Biolabs), with reagent volumes reduced to one-fifteenth of recommended volumes. For samples preserved in 80% ethanol, libraries were prepared without the enzymatic fragmentation step as the DNA of these samples is typically highly fragmented.

Libraries were pooled in equimolar concentration and paired-end sequenced using the Illumina HiSeq X or Novaseq platforms at a read length of 150 bp.

### *UCE loci identification and* in silico *bait design*

The identification of UCE loci was carried out using PHYLUCE *v*1.6.6 (Faircloth, 2016) following the recommendations of the tutorial (https://phyluce.readthedocs.io/en/stable/) and outlined workflow (Faircloth, 2017). Four publicly available genomes belonging to distantly related termite species were used to design baits: *Zootermopsis nevadensis* (Archotermopsidae), *Cryptotermes secundus* (Kalotermitidae), *Coptotermes formosanus* (Rhinotermitidae), and *Macrotermes natalensis* (Termitidae). Genome completeness was assessed using BUSCO *v*4.1.2 (Simão *et al*., 2015) and QUAST *v*5.0.2 (Gurevich *et al*., 2013). The genome of *M. natalensis* was chosen as the base genome for bait design due to its comparatively higher quality (for details, see Table S1).

Repetitive elements, retroelements, transposons, and small RNAs were masked from genome assemblies using RepeatMasker *v*4.1.1 (Smit *et al*., 2015) with the command line “-species eukaryota -div 50”. Assemblies were converted in the 2-bit format using the faToTwoBit tool of the BLAT suite of programs (Kent, 2002). We simulated 100 bp error-free paired-end sequencing reads from the three genome assemblies other than that of *M. natalensis* using art_illumina Q *v*2.5.8 (Huang *et al*., 2012) with the command line “--fcov 2 --mflen 200 --sdev 150”. In order to identify orthologous loci representing putative UCEs, the reads simulated from the three termite genome assemblies were mapped independently on the genome assembly of *M. natalensis* with a 0.05 substitution rate onto the base genome using the permissive raw-read aligner Stampy *v*1.0.32 (Lunter & Goodson, 2011). The three alignment maps were handled with SAMtools *v*1.9 (Li *et al*., 2009) and converted into BED files with bedtools *v*2.29.2 (Quinlan & Hall, 2010). In each BED file, putative conserved regions overlapping by at least one nucleotide were merged using bedtools. Conserved sequences shorter than 80 bp or containing over 25% of masked nucleotides were discarded using the phyluce program phyluce_probe_strip_masked_loci_from_set. The putative orthologous loci found across the four termite genomes were combined into a database using phyluce_probe_get_multi_merge_table (Supplementary Data 1). A total of 175,535 loci shared by the four termite genomes were identified and extracted using phyluce_probe_query_multi_merge_table and phyluce_probe_get_genome_sequences_from_bed, respectively. Extracted UCE sequences shorter than 180 bp were buffered to 180 bp by including 5’ and 3’ flanking regions in equal amounts with phyluce_probe_get_genome_sequences_from_bed (Supplementary Data 2).

A preliminary set of 120 bp baits was designed from the base genome of *M. natalensis* using phyluce_probe_get_tiled_probes. Baits targeted a region of 180 bp and overlapped in its center by 60 bp (at 2X tiling density). UCEs with ambiguous base calls and GC-content above 70% or below 30% were discarded from the bait set. Duplicates, defined as sequences having 50% identity over half of their length, were also removed from the bait set using LASTZ (Harris, 2007) implemented in the programs phyluce_probe_easy_lastz and phyluce_probe_remove_duplicate_hits_from_probes_using_lastz. In order to further identify and remove non-specific baits, we aligned the bait set (Supplementary Data 3) to the four genomes with phyluce_probe_run_multiple_lastzs_sqlite using a minimum identity threshold of 80% and minimum coverage of 83%. Sequences shorter than 180 bp were buffered to 180 bp by including 5’ and 3’ flanking regions in equal amounts and extracted from the alignments using phyluce_probe_slice_sequence_from_genomes. The loci shared by the four termite genomes were identified using phyluce_probe_get_multi_fasta_table and phyluce_probe_query_multi_fasta_table (Supplementary Data 4). The final UCE bait set was designed with phyluce_probe_get_tiled_probe_from_multiple_inputs, and duplicates were removed using LASTZ as described above (397,910 baits targeting 50,616 loci; Supplementary Data 5). This final set of loci was tentatively annotated using the GFF file (NCBI Annotation Release 100) from the *Z. nevadensis* genome assembly (GCF_000696155).

### Genome assembling and mining of phylogenetic markers

Adapters and low-quality bases were trimmed from raw reads using fastp *v*0.20.1 (Chen *et al*., 2018), resulting in a total of 4.55 to 448.64 millions of paired-end reads per sample (for details, see Table S1). Trimmed reads were assembled using metaSPAdes *v*3.13 (Nurk *et al*., 2017). The quality and completeness of assemblies were assessed with QUAST and BUSCO (Table S1). Mitochondrial genome scaffolds were identified in metaSPAdes assemblies and annotated using MitoFinder *v*1.4 (Allio *et al*., 2020). Nuclear ribosomal RNA genes (5S, 5.8S, 18S, and 28S) were extracted from metaSPAdes assemblies using barrnap *v*0.9 (https://github.com/tseemann/barrnap). UCE loci were extracted from metaSPAdes assemblies using the final set of termite baits we designed and the PHYLUCE suite of programs with parameter values set as recommended in the tutorial and previously published studies (Faircloth *et al*., 2015; Faircloth, 2017; Quattrini *et al*., 2018). Briefly, baits were aligned to the metaSPAdes assemblies at a minimum similarity threshold of 50% with phyluce_probe_run_multiple_lastzs_sqlite. Sequences of the metaSPAdes assemblies matching baits were extracted with the flanking 200 bp situated at both the 5’ and 3’ ends using phyluce_probe_slice_sequence_from_genomes. Extracted sequences were mapped back to the baits using phyluce_assembly_match_contigs_to_probes with a minimum identity of 80% over 67% of bait length to remove duplicates and sequences matching multiple UCE loci (Supplementary Data 6; Contribution #1 to the Termite UCE Database available at: https://github.com/oist/TER-UCE-DB/). The average coverage of UCE loci per sample was obtained using the mapping workflow of PHYLUCE *v*1.7.1.

### Sequence alignment

The 13 mitochondrial protein-coding genes, two mitochondrial rRNA genes, 22 mitochondrial tRNA genes, four nuclear rRNA genes, and UCEs were aligned using MAFFT *v*7.305 (Katoh & Standley, 2013). For mitochondrial protein-coding genes, we translated DNA sequences into the corresponding amino acid sequences using the transeq function from EMBOSS *v*6.6.0 (Rice *et al*., 2000) and aligned protein sequences with MAFFT. Protein alignments were back-translated into codon alignments using PAL2NAL *v*14 (Suyama *et al*., 2006). The other four types of genes, the mitochondrial rRNA and tRNA genes, nuclear rRNA genes, and UCEs, were aligned as DNA sequences. UCE loci were aligned using MAFFT implemented in phyluce_align_seqcap_align, and internal trimming was performed under default parameters with Gblocks (Castresana, 2000; Talavera & Castresana, 2007) implemented in phyluce_align_get_gblocks_trimmed_alignments_from_untrimmed. Loci absent in more than 25% of taxa were filtered out with phyluce_align_get_only_loci_with_min_taxa. The final UCE supermatrix was exported using phyluce_align_format_nexus_files_for_raxml (Supplementary Data 7: alignments; Supplementary Data 8: corresponding reduced bait set). Mitochondrial and nuclear gene alignments were concatenated using FASconCAT-G_v1.04.pl (Kück & Longo, 2014).

### Phylogenetic analyses

We ran one separate phylogenetic analysis for the mitochondrial genome alignment, the nuclear rRNA alignment, and the UCE alignment. We also ran one phylogenetic analysis for the combined UCE and mitochondrial genome alignments. The mitochondrial genome alignment was separated into five distinct partitions: combined rRNAs, combined tRNAs, and combined first, second, and third codon positions of protein-coding genes. Nuclear rRNA gene and UCE alignments were given a single partition each. Phylogenetic trees were reconstructed in a maximum likelihood (ML) framework using IQ-TREE *v*1.6.12 with 1,000 ultrafast bootstrap replicates (UFB) to assess branch supports (Nguyen *et al*., 2015; Chernomor *et al*., 2016; Hoang *et al*., 2018). The best-fit nucleotide substitution model was selected for each partition with the Bayesian Information Criterion using ModelFinder implemented in IQ-TREE (Kalyaanamoorthy *et al*., 2017). We calculated a global bootstrap support (GBS) value for each tree by averaging bootstrap values of all nodes. To assess concordance among UCEs, we carried out a multi-gene coalescence analysis with ASTRAL-III *v*5.7.7 (Zhang *et al*., 2018) using individual gene trees obtained with IQ-TREE as described above. We allowed polytomies to reduce gene tree biases. Branch supports calculated with ASTRAL represent local posterior probabilities (LPP), which are based on gene tree quartet frequencies (Sayyari & Mirarab, 2016). Topological conflicts between individual gene trees and the ASTRAL species tree were assessed with PhyParts (Smith *et al*., 2015) and visualized with PhyPartsPieCharts (https://github.com/mossmatters/phyloscripts/tree/master/phypartspiecharts).

## 3. Results

### In silico *data mining*

The mitochondrial genomes were retrieved from all 42 termite metaSPAdes assemblies. We also retrieved the four nuclear rRNA genes from 84% of the samples (see Table S1).

Our termite UCE bait set targeted a total of 50,616 loci distributed across 1,094 scaffolds of the *Z. nevadensis* genome assembly (GCF_000696155). Of these 50,616 loci, 6,325 (12.5 %) were found in the non-coding regions of 787 scaffolds (Supplementary Data 9). The remaining 44,291 loci were found in the coding regions (33,932 in exons) of 8,636 annotated genes distributed across 886 scaffolds. Of the 8,636 genes, 6,538 (75.7 %) contained more than one ultraconserved loci.

From the 50,616 targeted loci, we extracted between 3,426 and 42,860 non-duplicated UCE loci from 42 termite metaSPAdes assemblies (Table S1). The number of non-duplicated UCE loci extracted from the assemblies of *Cryptocercus* roaches varied between 13,480 and 16,331. The average coverage of UCE loci per sample was between 8.38 to 134.71x (Table S1). The final supermatrix, complete at 75% and containing loci present in at least 33 of the 45 taxa, was composed of 5,934 loci spanning over 1,677,394 nucleotide positions, 591,343 of which were parsimony-informative. The 45 taxa were represented by 939 to 5,928 loci.

### Phylogenetic reconstructions

Many deep and shallow relationships within termites were poorly resolved by the nuclear rRNA phylogenetic tree (GBS = 72.12) (Figure S1). Because of its poor performance, we excluded the rRNA alignment from the analysis performed on combined marker classes. The phylogenetic reconstruction based on mitochondrial genomes resolved most relationships (GBS = 87), except for several nodes within the Serritermitidae, the Rhinotermitidae, and the Termitinae (Figure S2), as previously reported (Bourguignon *et al*., 2015). The phylogenetic analysis performed exclusively on UCEs provided the most robust phylogenetic tree among the analyses performed on separate marker classes (Figure S3; GBS = 98.59). Combining UCEs and mitochondrial genomes marginally improved the resolution of the phylogenetic reconstruction (Figure 1; Figure S4; GBS = 99.02). The combined phylogenetic reconstruction resolved all nodes with high supports, except for the position of the rhinotermitid *Termitogeton planus* (UFB = 52 and 65, respectively). The phylogenetic analysis with ASTRAL revealed minimal discordance among the 5,934 UCE markers (Figure S5; final normalized quartet score of 0.89) (LPP = 1), except for five of the 42 nodes that presented conflicts among UCE markers. Within the Rhinotermitidae, the nodes corresponding to the split of *T. planus* and *Prorhinotermes simplex* displayed moderate concordance among UCE markers (LPP of 0.89 and 0.83, respectively). Within the Termitidae, the nodes corresponding to the split of *Neocapritermes utiariti, Pericapritermes* sp. 4, and *Nitiditermes* + *Cavitermes* showed moderate to high levels of discordance (LPP of 0.66, 0.98, and 0.39, respectively). PhyParts analyses on a subset of 1,000 gene trees revealed some levels of topological discordances (Figure S6). Nodes with discordance were mostly dominated by a plethora of topologies rather than by a single alternative and uninformative gene trees.

## 4. Discussion

We reconstructed phylogenetic trees for 42 species of termites and three species of *Cryptocercus* using three classes of markers: nuclear rRNA genes, mitochondrial genomes, and UCEs. The performance of the three types of phylogenetic markers decreased along the sequence: UCEs, mitochondrial genomes, and nuclear rRNA genes. The phylogenetic tree inferred from the latter class of markers, the nuclear rRNA genes, was poorly resolved and did not recover well-established relationships, such as the sister position of *Mastotermes* in respect to all other termites. The phylogenetic tree inferred from mitochondrial genomes was robust but failed to retrieve Sphaerotermitinae as sister to Macrotermitinae, as previously reported (Bourguignon *et al*., 2015; Bucek *et al*., 2019). The best phylogenetic tree was that reconstructed using the 75%-occupancy matrix comprised of 5,934 UCE loci (Figure 1). This phylogenetic tree was almost fully resolved and largely congruent with the phylogenetic trees inferred from transcriptomic data (Bucek *et al*., 2019). Therefore, our results indicate that the termite UCE bait set we designed performs very well when reconstructing phylogenetic relationships among termite species. The addition of mitochondrial genome data, which, as UCEs, can be recovered from shotgun genome assemblies, slightly improved the global bootstrap support of the termite phylogenetic tree.

The analysis with ASTRAL revealed a few cases of discordance among UCE markers for lineages of Rhinotermitidae and Termitidae whose phylogenetic position was also unresolved with transcriptomic data (Bucek *et al*., 2019). We used 5,934 UCE loci, a large number of markers that inevitably led to topological discordances between individual UCE trees and the species tree. These discordances are possibly caused by the lack of phylogenetic signal present in a single UCE marker and by population-level processes, such as incomplete lineage sorting and introgression, which frequently occurs during the emergence of new lineages (Degnan & Rosenberg, 2009; Blom *et al*., 2017; Parins-Fukuchi *et al*., 2021). While recent studies indicated that the phylogenetic resolution can be improved by removing loci localized in the exons of a single gene (Hedin *et al*., 2019; Van Dam *et al*., 2021), our unfiltered analysis performed as well as analyses performed on transcriptome-based data. The actual relationships among termite lineages with unresolved positions remain unclear, possibly reflecting intricate evolutionary history that cannot be satisfactorily resolved by molecular phylogenetic techniques.

We ran our analyses on samples for which we generated low coverage genome assemblies. The final bait set targeting a total of 50,616 orthologous loci was obtained from four termite genomes, belonging to four families. This high number of UCE loci is comparable to that found in other groups of arthropods (e.g., Buenaventura *et al*., 2021). Overall, 67% of loci in the termite bait set were found in exonic regions, further indicating that the UCEs of arthropods are mostly found in exons (Hedin *et al*., 2019; Van Dam *et al*., 2021). We retrieved numerous UCE sequences for all samples, including many that produced highly fragmented assemblies with low BUSCO scores (for details, see Table S1). All samples were accurately placed on the phylogenetic tree reconstructed with the 5,934 loci present in the 75%-occupancy supermatrix. Therefore, our UCE bait set has the potential to be used for mining phylogenetically informative genetic data from assemblies obtained from shotgun sequencing experiments. We established a centralized termite UCE database (https://github.com/oist/TER-UCE-DB/), which we plan to use to reference all UCE data extracted with the presently designed bait set, thereby ensuring the long-term re-usability of the available data.

While producing low-coverage genomes is more costly than targeting UCEs through synthetized baits, low-coverage genome data can be used to investigate a broad range of questions in addition to phylogenetic reconstruction. Used in combination with non-destructive DNA extraction protocols, our UCE baits could also be used to obtain sequence data from material that cannot be damaged, such as specimens from type series. This approach was successfully applied to centuries-old museum specimens of Opiliones, carpenter bees, and weevils (Blaimer *et al*., 2016; Van Dam *et al*., 2017; Derkarabetian *et al*., 2019). We recently obtained the full mitogenome of a Syntype of the termite *Archotermopsis wroughtoni* collected at the end of the 19^th^ century using shotgun sequencing data (Wang *et al*., 2021). Termite UCEs could be extracted using the same procedures. Termite taxonomy, which is led by a shrinking pool of experts and is largely based on soldier and worker gut morphology, could benefit from the use of the many UCE markers designed in this study (Eggleton, 1999; Korb *et al*., 2019). UCE baiting from whole-genome shotgun sequencing is the perfect tool to carry out a global taxonomic revision of termites.

## Supporting information

Main Electronic Supplementary Materials

## Acknowledgments

We thank Yves Roisin, Saran Traoré, and Guy Josens for providing specimens as well as Crystal Clitheroe for helping with DNA extraction and library preparation. We also thank the Scientific Computation and Data Analysis Section (SCDA) of the Okinawa Institute of Science and Technology Graduate University, Okinawa, Japan, for providing access to the OIST computing cluster.

## Notes

**Funding** This work was supported by the Czech Science Foundation (project No. 15-07015Y), the Internal Grant Agency of the Faculty of Tropical AgriSciences, CULS (20213112), the Japan Society for the Promotion of Science (JSPS) through a postdoctoral fellowship for foreign researchers awarded to SH (19F19819), and the Okinawa Institute of Science and Technology through core unit funding.

https://github.com/oist/TER-UCE-DB/

### Competing Interest Statement

The authors have declared no competing interest.

### Summary of Updates

Added coverage and annotation analyses Minor text editions

