## Supplementary material for "Using ultraconserved elements to reconstruct the termite tree of life": Main Electronic Supplementary Materials

**Running title**: UCEs in termites

Simon Hellemans[1,*], Menglin Wang[1], Nonno Hasegawa[1], Jan Šobotník[2], Rudolf H. Scheffrahn[3], Thomas Bourguignon[1,2,*]

[1]Okinawa Institute of Science & Technology Graduate University, 1919-1 Tancha, Onna-son, Okinawa 904-0495, Japan.

[2]Faculty of Tropical AgriScience, Czech University of Life Sciences, Kamýcka 129, 16521 Prague, Czech Republic.

[3]Fort Lauderdale Research and Education Center, Institute for Food and Agricultural Sciences, 3205 College Avenue, Davie, Florida 33314 USA.

### A. Supplementary Table

**Supplementary Table 1** (*See separate Excel file: Supplementary_File_2.xlsx*): (Sheet 1) Details of samples and external data used in this study. Publicly available genomes used for the UCE bait design are highlighted in grey. Assembly metrics were performed with QUAST *v*5.0.2 and are based on contigs of size ≥ 500 bp. BUSCO scores were obtained by screening against the insecta_odb10.2020-09-10, which tests for the presence of 1,367 single-copy ortholog groups. Abbreviations: C, complete BUSCOs; D, complete and duplicated BUSCOs; F, fragmented BUSCOs; M, missing BUSCOs; S, complete and single-copy BUSCOs.

### B. Supplementary Figures (*See separate PDF file: Supplementary_File_3.pdf*)

**Supplementary Figure S1**: Maximum likelihood reconstruction with IQ-TREE of the main termite lineages using the nuclear ribosomal RNAs (concatenated 5S, 5.8S, 18S and 28S) partition. Branch lengths are proportional to the number of substitutions per nucleotide site, and supports are ultrafast bootstrap values.

**Supplementary Figure S2**: Maximum likelihood reconstruction with IQ-TREE of the main termite lineages using the five mitochondrial partitions. Branch supports are ultrafast bootstrap values.

**Supplementary Figure S3**: Maximum likelihood reconstruction with IQ-TREE of the main termite lineages using the 75%-completeness (concatenated 5,934 UCE loci) partition. Branch lengths are proportional to the number of substitutions per nucleotide site, and supports are ultrafast bootstrap values.

**Supplementary Figure S4**: Maximum likelihood reconstruction with IQ-TREE of the main termite lineages using the combined 75%-completeness (concatenated 5,934 UCE loci) partition and mitochondrial partitions. Branch lengths are proportional to the number of substitutions per nucleotide site, and supports are ultrafast bootstrap values.

**Supplementary Figure S5**: Coalescent-based species tree produced by ASTRAL-III from the individual 5,934 single-gene maximum likelihood trees from IQ-TREE. Branch lengths are in coalescent units, and supports are local posterior probabilities.

**Supplementary Figure S6**: Topological conflicts between gene trees and the ASTRAL species tree as evidenced by PhyParts from a subset of 1,000 gene trees and visualized by PhyPartsPieCharts. Numbers indicate the number of concordant (above the branch) and discordant (below) gene trees. Pie charts reflect the proportion of gene trees with the concordant topology (in blue), the main alternative conflicting bipartition (in red), the alternative conflicting bipartition (in orange), and the remaining uninformative (supporting or conflicting) gene trees with <50% bootstrap (in grey).

### C. Supplementary Data (*will be submitted to Dryad*)

**Supplementary Data 1**: SQL database containing the putative orthologs found in the four termite genomes, obtained with ‘phyluce_probe_get_multi_merge_table’.

**Supplementary Data 2**: Sequences with equally distributed buffer, obtained with ‘phyluce_probe_get_genome_sequences_from_bed’, designed for the base genome of *M. natalensis*.

**Supplementary Data 3**: Deduplicated probe set (minimum identity threshold of 80% and minimum coverage of 83%) cleaned from sequences with ambiguous base calls and GC-content above 70% or below 30%, designed for the base genome of *M. natalensis* (240,602 baits targeting 170,648 loci).

**Supplementary Data 4**: Sequences with equally distributed buffer, obtained with ‘phyluce_probe_get_genome_sequences_from_bed’, designed for all four considered termite genomes (53,422 loci).

**Supplementary Data 5**: Final deduplicated probe set (minimum identity threshold of 80% and a minimum coverage of 83%) cleaned from sequences with ambiguous base calls and GC-content above 70% or below 30%, designed for all four considered termite genomes (397,910 baits targeting 50,616 loci).

**Supplementary Data 6**: Termite UCE Database. Extracted UCEs from all samples using the final deduplicated probe set (Supplementary Data 5) in which each sample was assigned a unique identification code (TER-X-UCEDB; see Supplementary Table 1). The database is maintained at: https://github.com/oist/TER-UCE-DB/.

**Supplementary Data 7**: Alignments produced for the 5,934 loci for which data missingness was below 25%. (A) Nexus alignment file. (B) Character sets file.

**Supplementary Data 8**: Bait set reduced to 5,934 loci (47,091 baits), corresponding to the loci kept in the 75% sample matrix (Supplementary Data 7).

**Supplementary Data 9**: Tentatively annotated UCEs, based on the GFF file (NCBI Annotation Release 100) from the *Z. nevadensis* genome assembly (GCF_000696155).
