## Supplementary Figures for "Using ultraconserved elements to reconstruct the termite tree of life"

**Figure S1. nuc rRNAs**

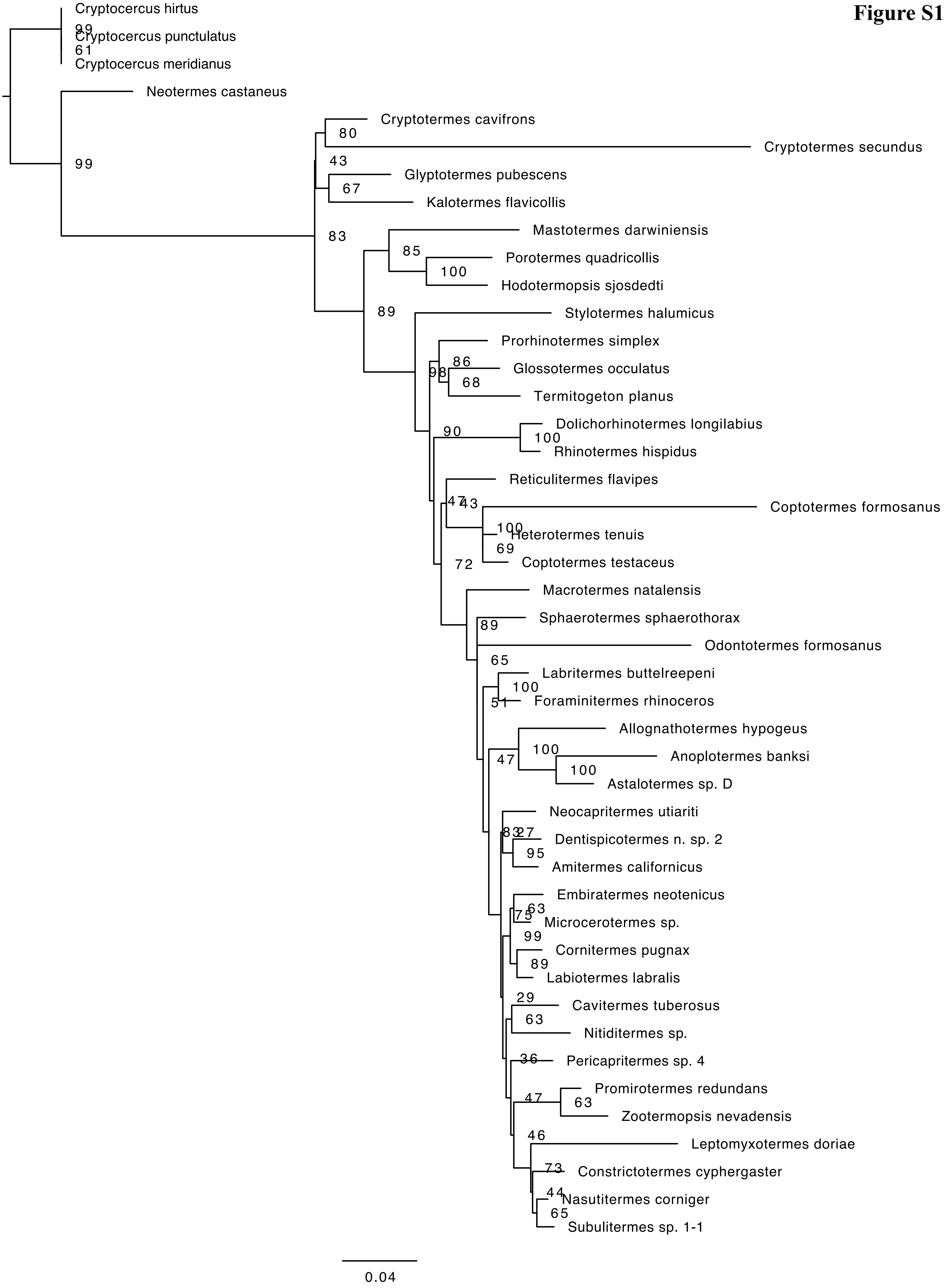

**Figure S2. Mitochondrial**

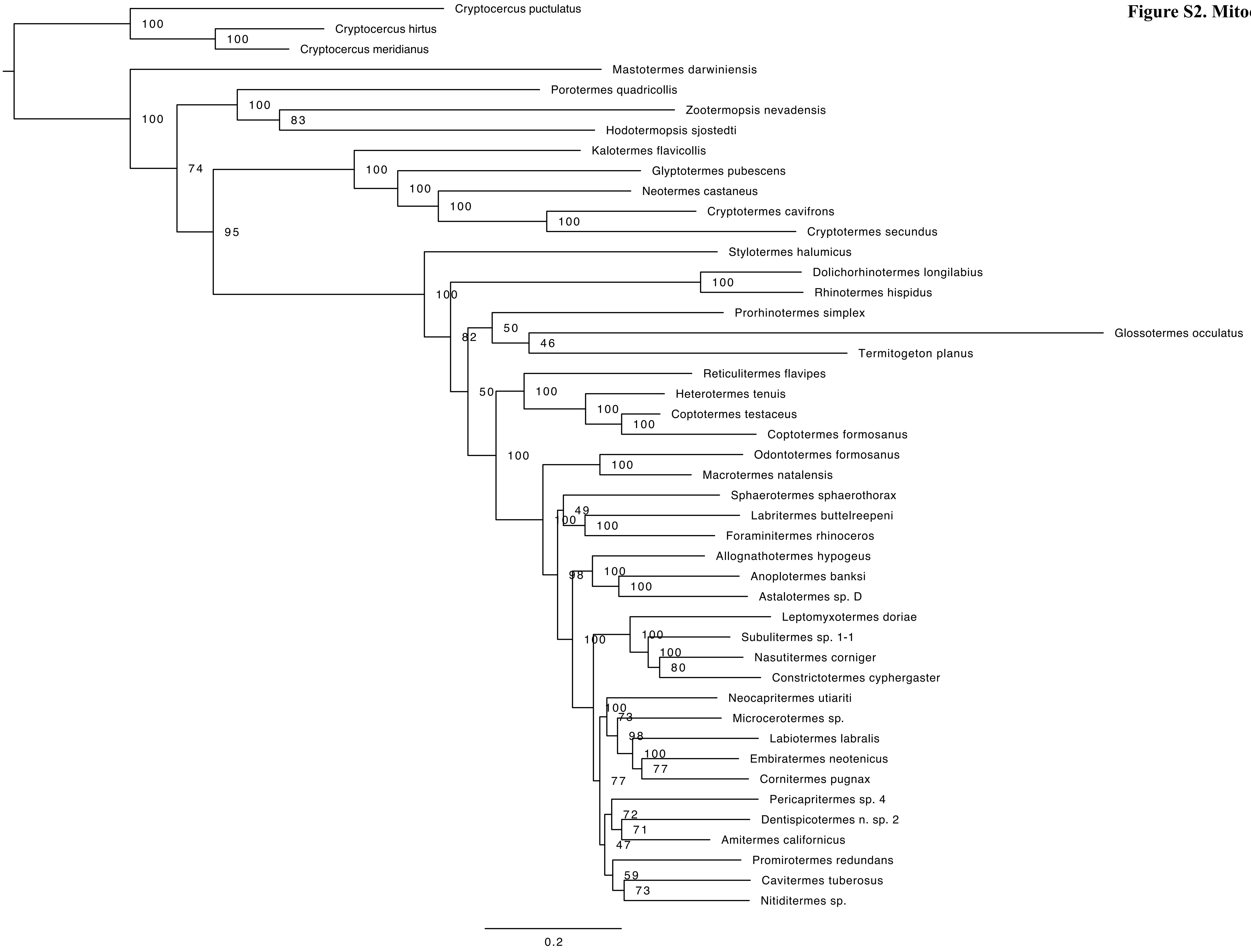

**Figure S3. UCEs supermatrix**

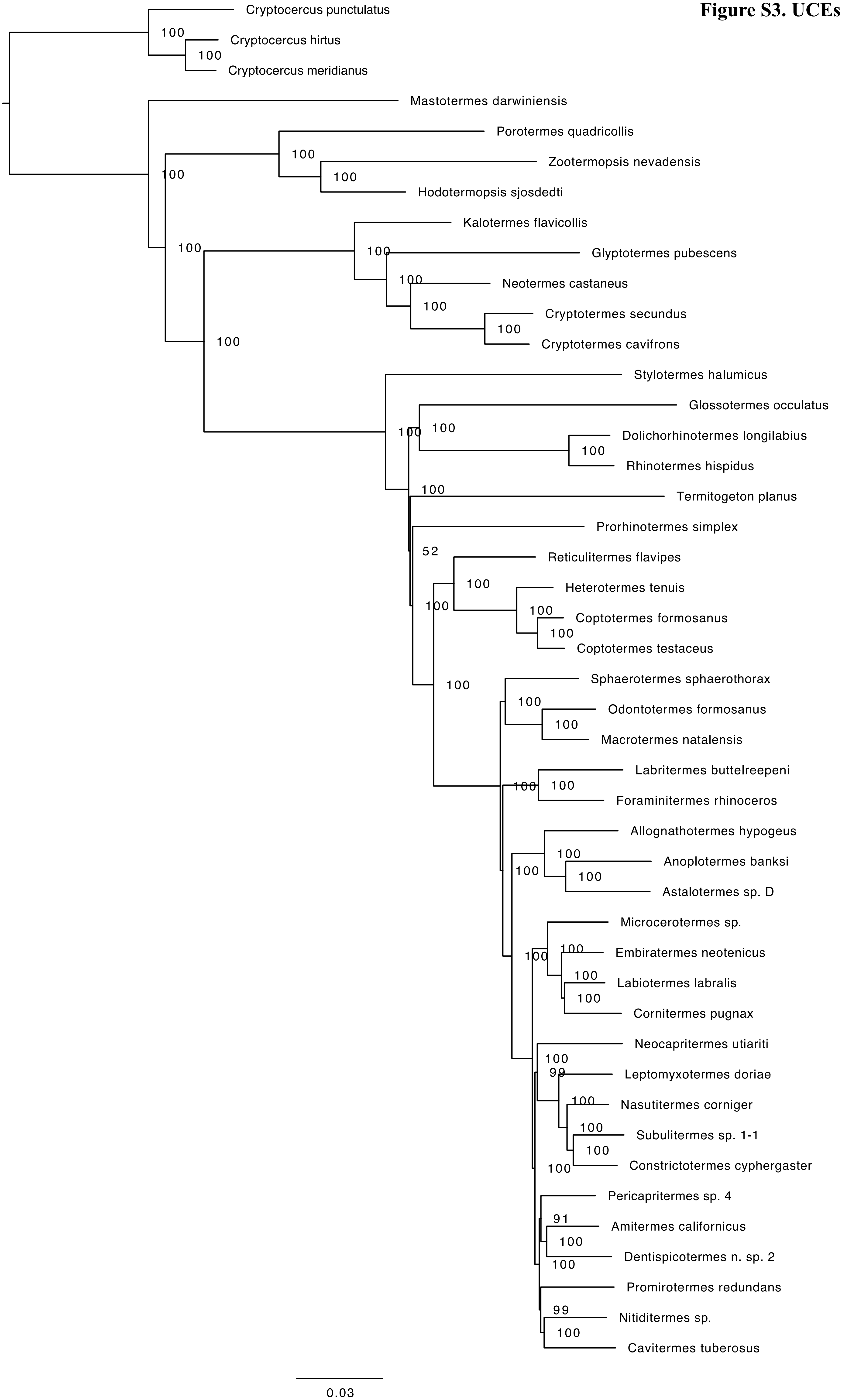

**Figure S4. combined UCEs & mitochondrial**

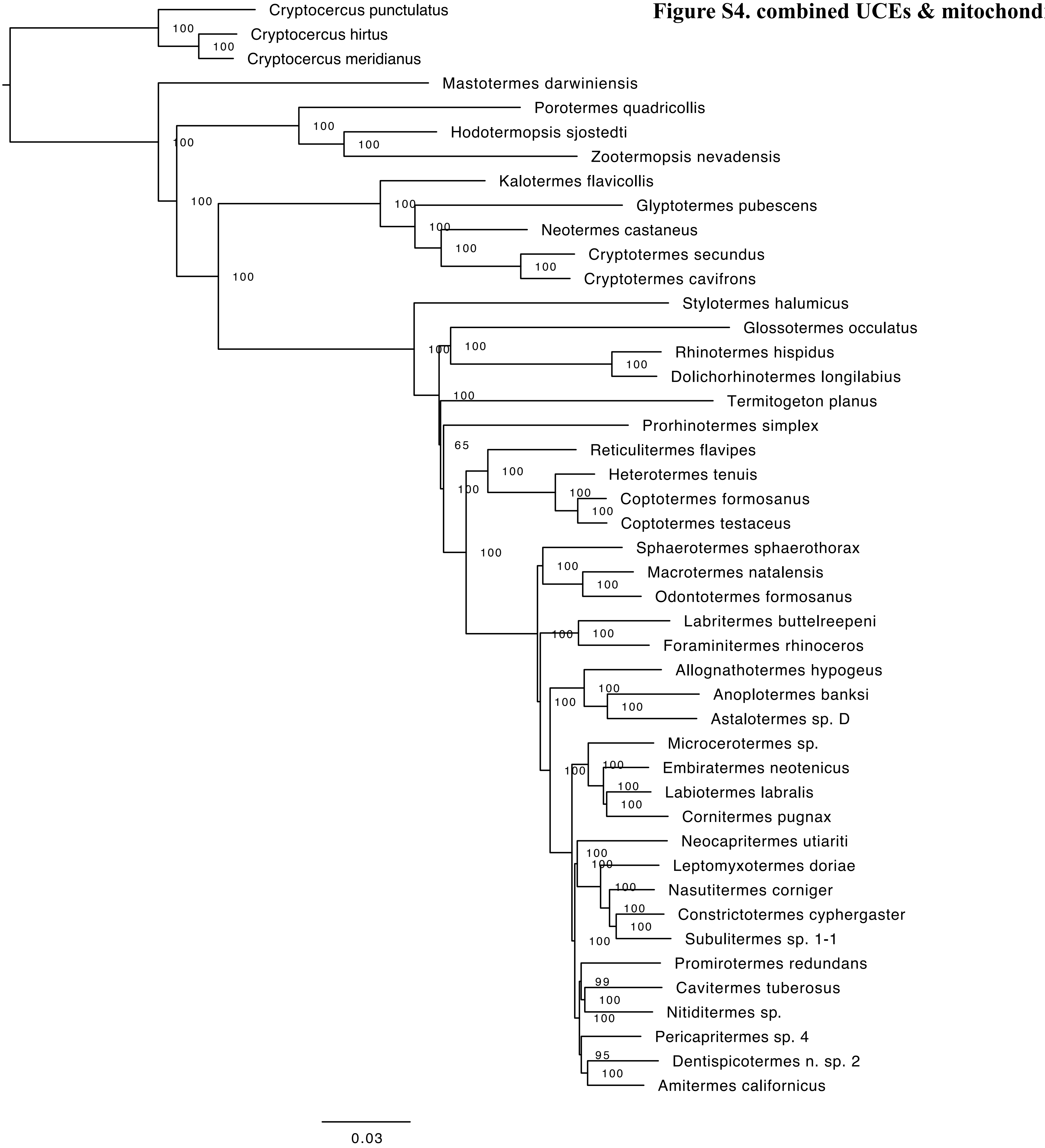

### Figure S5. ASTRAL UCEs

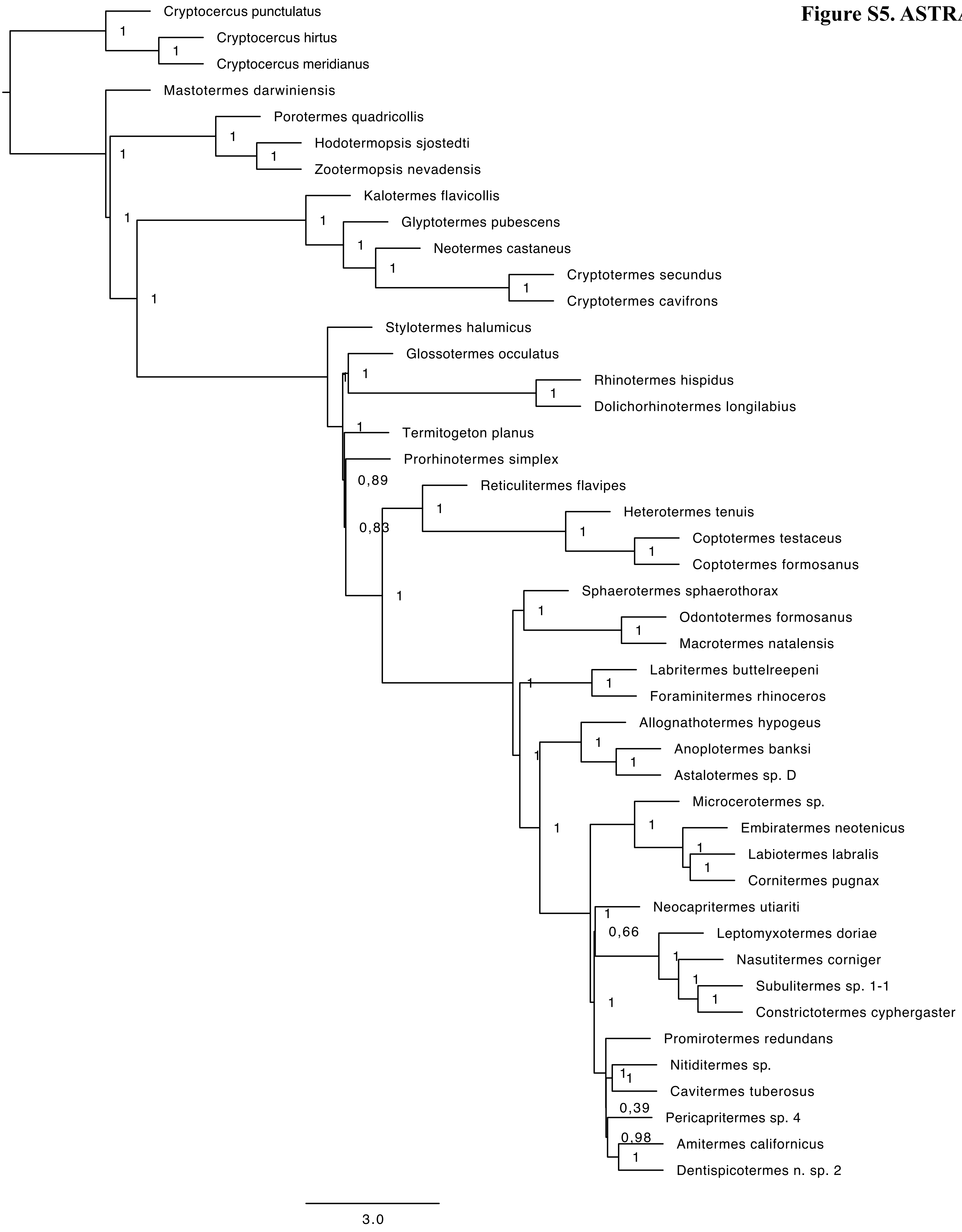

### Figure S6. PhyParts UCEs

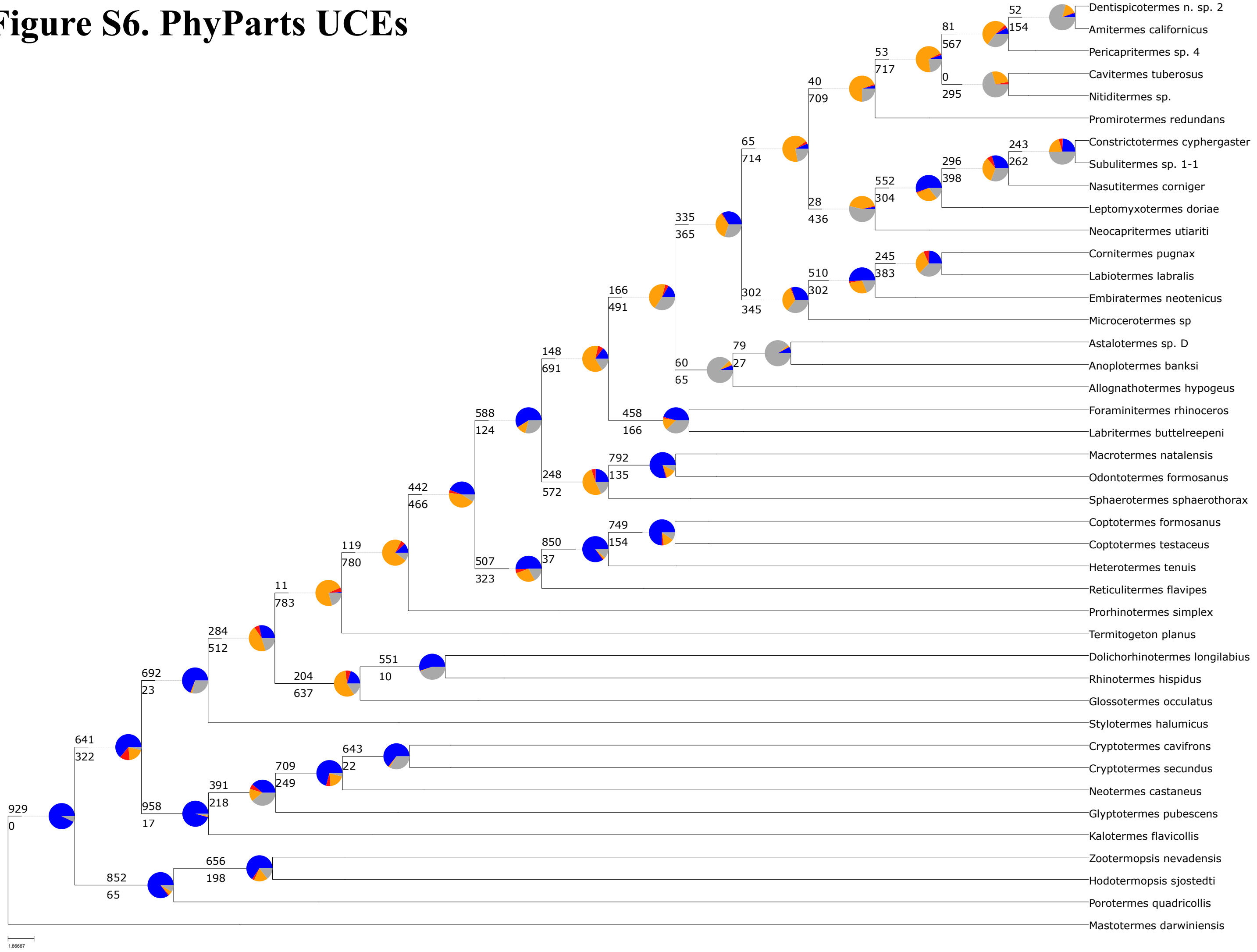
